# Discovering new amyloid-like peptides using all-atom simulations and artificial intelligence

**DOI:** 10.1101/2025.02.12.637911

**Authors:** Xiaohan Kuang, Sharareh Jalali, Tasnima Rahman, Jadhy Michalowski, Caren Sheng-Wong, Jirasak Wong-Ekkabut, Zhaoqian Su, Cristiano L. Dias

## Abstract

Establishing the fundamental relationships between peptide sequences and fibril formation is critical both for understanding protein misfolding processes and for guiding biomaterial design. Here, we combine all-atom molecular dynamics (MD) simulations with artificial intelligence (AI) to investigate how subtle variations in the arrangement of a short peptide sequence affect its propensity to form fibrils. Our results show that small shifts in the distribution of hydrophobic residues and charge clusters can significantly influence both the nucleation rate and the stability of cross-*β* structures. To rapidly extend this analysis over a wide sequence space, we developed an active learning–enhanced framework—Machine Learning for Molecular Dynamics (ML4MD)—that iteratively refines its predictions based on MD-derived aggregation data. ML4MD efficiently screens numerous peptide permutations and guides the discovery of previously unrecognized fibril-prone sequences, achieving an area under the receiver operating characteristic (ROC) curve (AUC) of 0.939. Overall, ML4MD streamlines the rational design of amyloid-like peptides by integrating detailed atomistic simulations with rapid and high-accuracy ML predictions.

## Introduction

Amyloid peptides are implicated in more than 30 degenerative diseases, including Alzheimer’s and Parkinson’s. ^1? ? –3^ They are known to aggregate in the form of fibrils that become deposited in different tissues gradually leading to organ failure. Despite their toxicity, fibrils exhibit mechanical properties that are desirable for various biomedical/biotechnology applications in addition to being biocompatible and biodegradable.^4–7^ Accordingly, non-toxic amyloids are being engineered for applications such as drug delivery systems, and antimicrobial gels.^8–10^ Bioinformatic tools informed by experimental studies have become an integral part of these design strategies. However, the accuracy of these in silico predictions has been questioned for sequences showing no degree of homology to existing amyloids. ^11–13^ This suggests that new methods are needed to *de novo* design sequences that have no resemblance to disease causing amyloids.

Amyloid fibrils are one-dimensional proteins aggregates that can grow up to several micrometers in length.^14–17^ At the molecular level, amyloid fibrils are characterized by multiple laminated *β*-sheets packed against each other to allow for cross-sheet side chain interactions known as the cross-*β* pattern.^18^ All proteins, independent of their sequence, are believed to form fibril-like structures at high concentration under the right solvent conditions. ^19^ However, the amino acid sequence strongly modulates this process enabling some proteins to form fibrils at physiological or ambient conditions while others require extreme conditions to do so. Amyloid proteins are usually small and found to be intrinsically disordered in dilute solution adopting intermittently small *β*-sheet elements along the main chain. ^20–23^ This structural complexity allows for polymorphism making the direct prediction of fibril structure from sequence particularly challenging.^24^ For instance, the predictive ability of AlphaFold2 for fibril structures is limited, often showing an unexpected preference for mated *α*-helical sheets. This limitation is likely due to the imbalanced training dataset, where the Protein Data Bank comprises 98% globular proteins, 2% transmembrane proteins, and almost no fibril structures. ^25^

Some bioinformatics tools to predict amyloidogenicity are based on determining short segments within the sequence that are hot spots for fibril formation. ^26^ These segments are determined based on the propensity of their amino acids to form *β*-sheets (TANGO^27^ and Zyggregator^28^) and/or fibrils (AGGRESCAN^29^ and FoldAmyloid^30^). Various physico-chemical properties of the different residues are also considered when predicting fibril formation as well as the flanking of hot-spot segments by charged amino acids of the same sign. ^28^ Since folding into a given structure is the result of competition between different conformations, some algorithms consider the propensity of a sequence to fold into non-*β*-sheet structures. ^27^ Others consider the ability of side chains to interact within neighboring *β*-strands (PASTA^31?^ and BETASCAN^32^) or they compute energies of peptides when folded into known cross-*β* structures.^33^ Experimental databases of peptides that form fibrils have also been compiled, ^27,34^ and they are used to test or as input for machine learning algorithms to make predictions. Existing tools to predict fibril formation from the amino acid sequence are knowledge based. Accordingly, they tend to be relatively accurate for sequences that show a high degree of homology to known amyloid peptides. However, for peptides that differ significantly from those found in compiled databases, predicted propensities are not always reliable.^11,13^ As an illustrative example, for sequences made of phenylalanine (F), lysine (K), and glutamic acid (E), TANGO and AGGRESCAN predictions do not match experimental measurements.^12,35^ Experimentally,^36^ three isomeric sequences composed of F, K, and E decrease in amyloid propensity in the following order: (FKFE)_2_ *>* KEFFFFKE *>* (KFFE)_2_. TANGO incorrectly predicts the reverse order of fibril formation for these sequences, while AGGRESCAN predicts the same propensity for (FKFE)_2_ and (KFFE)_2_. This is surprising as (KFFE)_2_ does not form fibrils experimentally even at high concentration (1.0 mM) whereas (FKFE)_2_ promptly forms fibrils at low concentration (0.2 mM). Clearly, existing algorithms are deficient based on our own lack of understanding regarding the fundamental rules for amyloid self-assembly.

Unbiased molecular dynamics (MD) simulations rely solely on our understanding of how atoms interact with each other and do not require prior knowledge of the system being studied. Thus, MD simulations could be accurate in describing fibril formation of sequences that show no degree of homology to known amyloids. The coarse grained Martini force field was used to measure aggregation in simulations of 400 di-peptides, ^37^ 8,000 tri-peptides, ^38^ and 160,000 tetra-peptides. ^39^ These studies show that peptides with a high propensity to aggregate have a preference for displaying positively and negatively charged amino acids at N- and C-terminals, respectively, with aromatic residues being preferentially located in the middle of sequences.^38^ One drawback of the Martini force field is that it does not consider hydrogen bond formation; therefore, it cannot be used to predict the formation of *β*-sheets and amyloid fibrils. Therefore, steric constraints that emerge from the proximity of side chains in fibril-like structures cannot be predicted using this coarse grain model. In addition, peptide-peptide interactions were significantly reduced in recent versions of the Martini force field enabling it to describe the behavior of proteins inside lipid membranes but disabling aggregation in solution. ^40^

Recently, all-atom models have been able to describe the formation of amyloid fibrils from unbiased initial conformations for short peptide sequences. ^12,35,41–43^ This is being used to describe the spontaneous nucleation of fibrils via primary and secondary mechanisms as well as their growth into longer fibrils.^44,45^ Moreover, the time required to nucleate fibrils in these simulations was shown to depend on the peptide sequence. For the three peptides made with F, K, and E amino acids, the nucleation time increases in the same order as in experiments, ^12^ i.e., (FKFE)_2_ *>* KEFFFFKE *>* (KFFE)_2_. This provides evidence that all-atom simulations may be used to predict the relative propensity of a peptide to form fibrils even if its sequence is significantly different from the ones found in amyloid databases. However, significant computational resources are needed for peptides to nucleate into fibrils in MD simulations. Moreover, several weeks are often required for these simulations to be completed while results may be needed within minutes during the design phase of new materials.

In this study, we introduce a novel framework that combines molecular dynamics (MD) simulations and machine learning (ML) to investigate how peptide sequences affect fibril formation. Our approach proceeds in several stages. First, we perform long-timescale MD simulations in large boxes to capture the nucleation and growth processes of amyloid-like fibrils across various peptide sequences, confirming that both hydrophobic/polar arrangement and charge clustering critically influence fibril-forming propensity. Next, to accelerate screening, we conduct shorter MD simulations in small, high-concentration boxes, demonstrating a fast and reliable method for evaluating early-stage aggregation. We compile the outcomes of these high-concentration simulations into a database, linking each peptide sequence to its observed fibril formation tendency. This database then serves as the training set for an ML model that can predict the fibril-forming potential of new peptide sequences. Finally, leveraging this trained model, we rapidly explore the broader sequence space and uncover additional motifs that favor fibril formation in mere seconds of computational time. These newly identified sequences are re-examined through MD simulations, annotated, and added to our training set in an active learning loop. By following this iterative strategy, we achieve an area under the ROC curve (AUC) of 0.939 (±0.07). We refer to this integrated methodology as **Machine Learning for Molecular Dynamics (ML4MD)**. By uniting detailed atomistic simulations with fast ML predictions, ML4MD offers a powerful tool for understanding amyloid formation and speeds the rational design of functional amyloid-like materials, as well as the development of anti-amyloid strategies.

## Results

### Molecular Dynamics

#### Simulating large boxes

Fig. 1 revisits the aggregation behavior of three peptides composed of permutations of four phenylalanines (F), two lysines (K), and two glutamic acids (E). In computer simulations performed at very high concentration (37 mM),^12^ these peptides with sequence Ac-(FKFE)_2_-NH_2_ (panel a), Ac-KEFFFFKE-NH_2_ (panel b), and Ac-(KFFE)_2_-NH_2_ (panel c) showed very different propensities to form fibrils. Specifically, Ac-(FKFE)_2_-NH_2_ rapidly formed fibrils, while Ac-KEFFFFKE-NH_2_ showed a slower fibrillation rate, and Ac-(KFFE)2-NH_2_ did not form fibrils under the conditions tested. A similar trend was also reported experimentally. ^36^ In Fig. 1, simulations are conducted at a peptide concentration of 7.5 mM, starting with ten peptides randomly positioned in solvated cubic box. These simulations with close to 200,000 atoms are performed for 3 microseconds, and the structure of the largest aggregate at the end of this period is illustrated in panels *a-c*. To assess aggregation, the size *S*_clus_ of the largest aggregate/cluster (shown in green) and the number 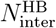 of inter-peptide hydrogen bonds (depicted in black) are plotted over time in panels *d-f*.

**Figure 1.**
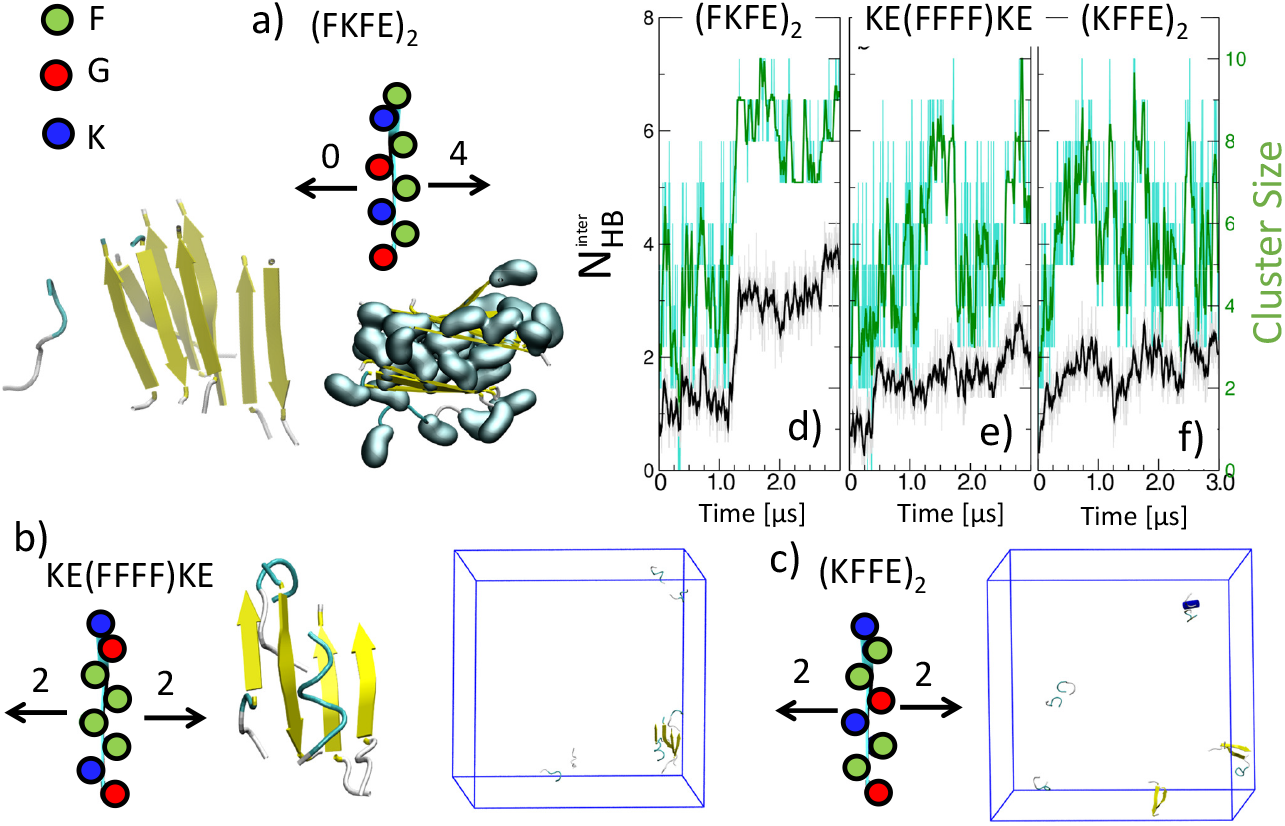
Revisiting simulations of three peptides that form fibrils with different propensities. Final configuration in simulations performed with peptides that have (a) non-polar and charged residues alternating along the sequence, (b) four consecutive non-polar residues flanked by pairs of charged residues at both ends, (c) two pairs of consecutive non-polar residues separated by a neutral pair of charged amino acids. Yellow arrows represent *β*-sheets and non-polar phenylalanine side chains are drawn using a cyan surface contour. (d-f) Number of inter-backbone hydrogen bonds (black lines) and largest cluster size (green lines).

Before the end of the simulation, Ac-(FKFE)_2_-NH_2_ peptides promptly form a cross-*β* structure with four non-polar residues buried in between the *β*-sheet bilayer–Fig. 1a. These fibril-like structures emerge abruptly in our simulations at time ∼ 1.25 *µ*s when *S*_clus_ and 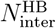 increase simultaneously–panel d. Peptides with sequence Ac-KE**F**_4_KE-NH_2_ aggregate more gradually, culminating in a single *β*-sheet made with four peptides at the end of the simulation–Figs. 1b,e. In contrast, Ac-(KFFE)_2_-NH_2_ peptides only formed dimers intermittently in the simulation, which did not progress to larger aggregates–Figs. 1c,f. Thus, our simulations indicate a decreasing propensity for fibril formation in the order: Ac-(FKFE)_2_-NH_2_ *>* Ac-KE**F**_4_KE-NH_2_ ≥ Ac-(KFFE)_2_-NH_2_, aligning with findings from previous studies. ^12,36^ Notice that values of *S*_clus_ and 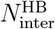 are not significantly different for Ac-KE**F**_4_KE-NH_2_ and Ac-(KFFE)_2_-NH_2_ peptides (panels e and f), likely due to the limited number of peptides involved in the simulations.

Our first hypothesis is that the relative position of non-polar residues along the sequence determines the ranking of peptides in terms of their propensity to form fibrils. This positioning defines how many non-polar side chains are oriented in the same direction when the peptides form *β*-strands, which is crucial for initiating and stabilizing cross-*β* structures. According to this hypothesis, the rate of fibril formation would be higher for Ac-(FKFE)_2_-peptides as it has four phenylalanine side chains facing the same direction (panel a), compared to Ac-KE**F**_4_KE-NH_2_ (panel b) and Ac-(KFFE)_2_-NH_2_ (panel c) peptides that have only two.

To test this hypothesis, additional simulations are carried out using the three peptide sequences depicted in Fig. 2. Two of these sequences, Ac-KK**F**_4_EE-NH_2_ (panel a) and Ac-**F**_4_KKEE-NH2 (panel b), have two non-polar side chains facing the same direction when peptides adopt a *β*-strand configuration. Surprisingly, these peptides rapidly aggregate into large clusters with *S*_clus_ values of 10 (panels *d, e*). This aggregation behavior contrasts with peptides in Fig.1*b,c* that have an equal number of non-polar residues oriented in the same direction but show only mild aggregation (Fig.1b) or no aggregation at all (Fig.1c). The third simulated peptide in Fig. 2c (with sequence Ac-KFFFKFEE-NH2) displays three non-polar residues facing the same direction when adopting a *β*-strand. Based on our hypothesis, the latter is expected to form larger and faster clusters compared to peptides in panels a,b. However, the opposite is observed: it forms smaller clusters in our simulations with *S*_clus_ values around 7. This disprove our initial hypothesis and underscore the existence of other patterns in the peptide sequence that may be as important as hydrophobic interactions in fibril formation.

**Figure 2.**
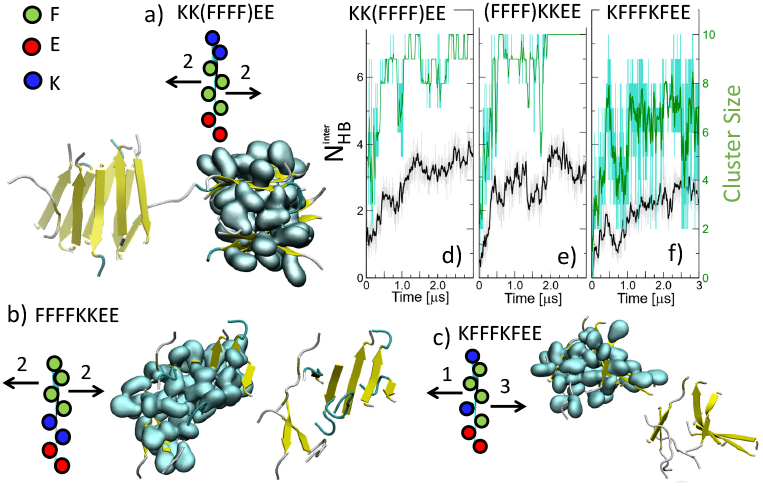
Simulations of three novel peptides made with 4 phenylalanines, 2 lysines, and 2 glutamic acids. Final configuration in simulations performed with peptides that have (a) four consecutive non-polar residues flanked by pairs of positive and negative residues at N- and C-termini, (b) four consecutive non-polar residues at the N-terminal, (c) three consecutive non-polar residues. Yellow arrows represent *β*-sheets and non-polar phenylalanine side chains are drawn using a cyan surface contour. (d-f) Number of inter-backbone hydrogen bonds (black lines) and largest cluster size (green lines).

Fig. 2a shows that Ac-KK**F**_4_EE-NH_2_ peptides form a bilayer consisting of two well-defined antiparallel *β*-sheets, with non-polar residues embedded between them. Albeit being more dis-order, *β*-sheets by Ac-KFFFKFEE-NH_2_ peptides are also packed against each other resembling fibril-like structures–Fig. 2c. These two sequences have oppositely charged N- and C-termini that may drive/facilitate packing and antiparallel alignment of *β*-sheets. In contrast, the lack of oppositely charged termini can result in the formation of less ordered *β*-sheets that are more difficult to pack against each other as observed in simulations of Ac-KE**F**_4_KE-NH_2_ (Fig.1b) and Ac-**F**_4_KKEE-NH_2_ (Fig. 2b) peptides. These observations suggest that in addition to hydrophobic interactions, electrostatics can also be crucial in influencing fibril formation. This highlights the complexity of predicting fibril formation based solely on peptide sequence, which requires a deeper understanding that can be obtained through the simulations of various peptide sequences and extensive analysis of these results.

#### Simulating small boxes

Simulations in Figs. 1–2 tracked a large number of particles (more than 200,000 atoms in order to simulate 10 peptides at concentrations of ∼ 7.5 mM) for many time-steps to account for 3 *µ*s. This type of simulation provides mechanistic insights into fibril formation.^12,44,45^ However, they are time consuming, which makes them impractical to explore a large set of peptide sequences. An example of the latter task involves screening all possible permutations made with 4 phenylalanines, 2 lysines, and 2 glutamic acids in order to identify nonpolar-polar sequence patterns that have a high tendency to form fibrils. This would involve simulating 420 sequences 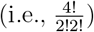, which is a demanding task for today’s computers.

To address fibril formation for a large set of sequences, *we hypothesize that interactions between peptides during the early stages of aggregation are predictive of a sequence’s propensity to form fibrils*. In light of this hypothesis, peptides that form fibrils would have a high tendency to interact via inter-peptide Hbonds (as opposed to intra-Hbonds) adopting *β*-structures when close to each other in space. To test this hypothesis, we perform simulations at a peptide concentration of 370 mM, i.e., 8 peptides randomly embedded in solvated cubic boxes of size 6 nm. At these high concentrations, peptides spend most of their time interacting with each other as opposed to diffusing in solution. These systems (composed of ∼ 20,000 atoms) can be simulated with much less resources than the ones in Figs. 1–2. Furthermore, since we are only interested in the early stages of aggregation, simulations can also be performed for shorter periods of time. Here, we simulate systems for 0.5 *µ*s.

Fig. 3 shows results from simulations performed using the six reference sequences in Fig. 1–2. The latter figure depicts the average number N_*β*_ of residues that adopt *β*-structures, and the difference ΔHbonds between average inter- and intra-backbone hydrogen bonds. Simulations in which peptides form fibrils are characterized by a large numbers of N_*β*_ and ΔHbonds. For each sequence, averages are computed from the last 0.1 *µ*s of four independent simulations differing from each other by the initial position of peptides in the simulation box and initial atomic velocities. Fig. 3 highlights the stochastic nature of the aggregation process where the computed quantities can vary between the different independent simulations. However, a clear trend is observed by comparing the set of independent simulations across different sequences. In particular, all simulations using Ac-(KFFE)_2_-NH_2_ (orange) peptides exhibit low numbers of N_*β*_ and ΔHbonds; whereas most trajectories performed with Ac-(FKFE)_2_-NH_2_ (black), Ac-(KF_3_KFEE)-NH_2_ (brown), Ac-F_4_KKEE-NH_2_ (green), and Ac-KKF_4_EE-NH_2_ (blue) result in high numbers for N_*β*_ and ΔHbonds. Notice that latter and former sequences correspond to peptides that remained dispersed in the solvent and aggregated rapidly in Figs. 1–2, respectively. Only simulations with Ac-KEF_4_KE-NH_2_ (red) peptides account for an equal number of trajectories with high and low values of N_*β*_ and ΔHbonds.

**Figure 3.**
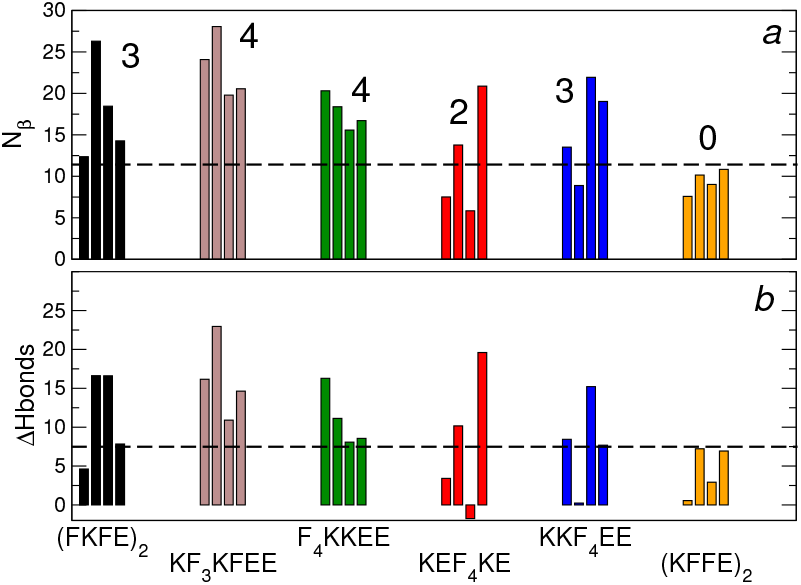
Average quantities computed from MD simulations performed in small boxes using the six reference sequences studied in Figs. 1–2. Four independent simulations are performed for each sequence and the average number of (a) residues that form *β*-structures, and (b) the difference ΔHbond between inter- and intra-backbone hydrogen bonds are shown. Dashed horizontal lines correspond to N_*β*_ = 12 and ΔHbonds = 7. The numbers of fibril-forming trajectories for each sequence are depicted.

To quantify the relative propensity of a sequence to form fibrils, we compute the number of simulations in which peptides have formed more than some given cutoff values of N_*β*_ and ΔHbonds. Trajectories that satisfy these conditions are called *fibril-forming trajectories*. These cutoffs are chosen empirically such as to allow for a high number of fibril-forming trajectories for sequences Ac-(**F**K**F**E)_2_-NH_2_, Ac-KFFFKFEE-NH_2_, Ac-**F**_4_KKEE-NH_2_, and Ac-KK**F**_4_EE-NH_2_; an intermediate number for sequence Ac-KE**F**_4_KE-NH_2_; and a low number for Ac-(K**FF**E)_2_-NH_2_–as expected from simulations in Figs. 1–2. Empirically, we find that this classification can be achieved with the cut-off values 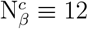 and ΔHbonds ^c^ ≡ 7. These cutoff values are shown as dashed lines in Figs. 3.

Although there is some subjectivity in this definition, we find that the trend in the relative number of fibril-forming trajectories is maintained for different sequences for reasonable variations in the cut-offs. For each of our reference sequences, the number of fibril-prone trajectories is indicated in Fig. 3.

#### Simulating many sequences

Our goal is to simulate a large number of peptides to provide an understanding of how the propensity to form fibrils can be predicted from the amino acid sequence. Out of all the 420 sequences that can be created via the permutation of 4 phenylalanines, 2 lysines, and 2 glutamic acids, 88 of them (including the six reference sequences of Fig. 1–2) are simulated using our small box protocol. These 88 peptides were chosen such that their sequences sample all 4-amino acid segments, e.g., FFFF or FKFE. Each sequence is made up of five 4-amino acid segments and there are sixty-three possible segments in the 420 peptides. The selected 88 peptides account for at least 20% of the possible occurrence of the 63 segments. Four independent simulations were conducted for each peptides and Fig. 4 shows the correlations between ΔHbonds and N_*β*_ computed for all the 88 × 4 trajectories. In this figure, dashed lines represent the cut-offs ΔHbonds^c^ ≡ 7 and 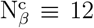 used to define fibril-forming trajectories.

**Figure 4.**
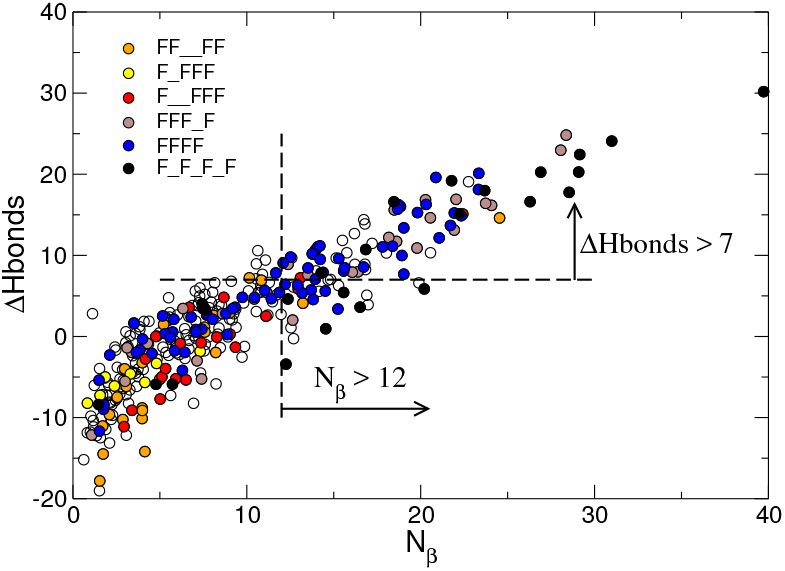
Correlation between ΔHbonds and N_*β*_ for all 4 × 88 simulated trajectories. Six polar-nonpolar sequence patterns are highlighted using different colors. Dashed lines represent the cutoffs used to define fibril-forming trajectories.

The color scheme in this figure provides insight into the role played by the polar-nonpolar sequence pattern in fibril formation. In particular, sequences where nonpolar and polar residues strictly alternate (`F_F_F_F,` black points), and sequences where three consecutive nonpolar residues are followed by polar and nonpolar amino acids (`FFF_F`, brown) have probabilities of 0.54 and 0.57 to form fibrils. These high probabilities may be due to the ability of these peptides to segregate nonpolar amino acids to one face of a *β*-sheet, which facilitates their burial away from the solvent when they form fibril-like structures. The trajectory of sequences with four consecutive nonpolar residues (`FFFF`, blue), have a slightly smaller probability (0.42) to form fibrils. In contrast, sequences where nonpolar residues are separated from each other by two charged residues (`FF__FF`, orange, and `F__FFF`, red), have very low probabilities to form fibrils–0.04 and 0.1, respectively. Notice that these sequences can segregate polar and nonpolar residues to different faces of helical structures and, thus, they may adopt non-extended conformations more frequently precluding them from forming fibrils. Fig. 4 also depicts two peptides containing one nonpolar residue followed by a charged and three nonpolar residues– `F_FFF`, yellow. None of the 8 trajectories of these peptides formed fibrils, albeit their non-polar pattern is the reverse (i.e., can be read backward) of `FFF_F` (brown), which forms fibrils with high probability. This suggests that symmetry may not be used to reduce the sequence space to be sampled.

In addition to the polar-nonpolar pattern, the relative position of negative and positive amino acids also affects fibril formation. For a given polar-nonpolar pattern, there are six permutations of two lysines and two glutamic acids, i.e., 4!/(2 × 2). The latter permutations for five polar-nonpolar patterns are shown in Fig. 5. Significant differences in the numbers of fibril-forming trajectories can be observed within the same polar-nonpolar pattern. For example, the number of fibril-forming trajectory changes from 3 to 0 by interleaving “E, K, K, E” into `F_F_F_F_` instead of “K, E, K, E”–fourth and first rows in the first column of Fig. 5. Thus, being able to segregate polar and non-polar residues to different faces of a *β*-strand does not guarantee that a peptide will necessarily form a *β*-sheet/fibril. Conversely, the same sequence of positive and negative residues can yield very different results when combined with different polar-nonpolar patterns. An example of the latter are the two peptides made by alternating polar and nonpolar residues (`F_F_F_F_` and `_F_F_F_F`) wherein the sequence of polar residues is “E, K, K, E”– fourth row of the first two columns in Fig. 5. Numbers of fibril-forming trajectories for these peptides are 0 and 3, respectively. Similar behaviors can be observed for peptides made using four consecutive non-polar residues located at the beginning, end, and middle of the sequence– blue cells in Fig. 5.

**Figure 5.**
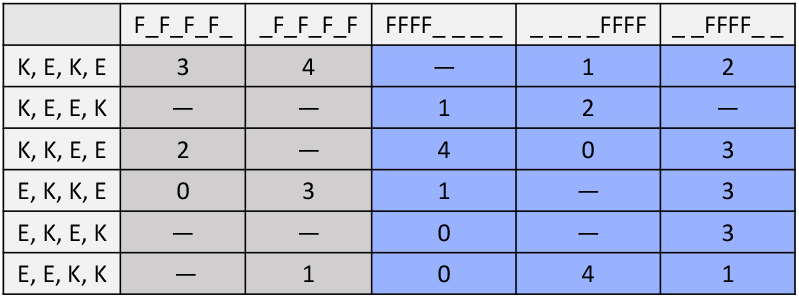
Number of fibril-prone trajectories for selected peptide sequences. Each column corresponds to a different polar-nonpolar pattern, and each line represents one of the six permutations of 2 lysines and 2 glutamic acids. Gray–Cells representing sequences with alternating polar and non-polar residues. Blue– Cells representing sequences made with four consecutive non-polar residues.

We anticipate that the sequence of positive and negative amino acids can affect fibril formation by promoting secondary structures in which the backbone of the peptide is not extended (e.g., *α*-helix or turn) thus inhibiting the formation of *β*-sheets and fibrils. In addition, positive and negative amino acids can also affect the structure and stability of *β*-sheets specially when buried inside the dry core of fibrils.^46^ Instead of trying to unravel these intricacies for all polar-nonpolar patterns, we turn to machine learning in order to predict fibril formation.

## Machine Learning

Building upon the molecular dynamics simulations dataset, we develop a machine learning (ML) model to predict the probability of a peptide to form fibrils. For simplicity, we refer to the “number of fibril-forming trajectories” computed from simulations as the ‘MD score’. For the purpose of training the model, sequences with MD scores of 2, 3, and 4 are classified as “fibril” (positive samples), whereas those with MD scores of 0 and 1 are classified as “non-fibril” (negative samples). Sequences scoring 1 were excluded from the training set due to their potentially ambiguous nature.

In our ML model, each amino acid is represented by 320 features derived from Evolutionary Scale Modeling (ESM-2) embeddings.^47^ The latter was trained to predict properties of randomly masked amino acids within protein sequences. Consequently, the local and global sequence context are encoded within ESM-2 embeddings, which can be used to inform about the structure and function of proteins/peptides. ^48,49^ In the ML model, we represent each sequence by the 320 features obtained by averaging over the eight residues. The goal of the model is then to predict how these 320 average features account for positive (fibril) and negative (non-fibril) samples.

The ML model consists of a Multilayer Perceptron (MLP) classifier with an architecture that includes an input layer with 320 neurons activated by a Rectified Linear Unit (ReLU) function, followed by four hidden layers with 256, 128, 64, and 32 neurons, respectively, each with ReLU activation. To prevent overfitting, each hidden layer is followed by a dropout layer with a rate of 0.2. L2 regularization (coefficient: 0.01) is applied to each hidden layer to mitigate overfitting. The output layer is a single neuron with sigmoid activation for binary classification, determining whether a given sequence is fibril-forming. The model is trained using binary cross-entropy loss and optimized with the Adam optimizer. Early stopping, with a patience of 20 epochs, is employed to prevent overtraining, and the best model weights are restored.

An active learning framework is employed to iteratively train the ML model in order to efficiently explore the entire sequence space (Figure 6). This process involves a cycle of training, sequence prediction, and re-evaluation, allowing for continuous model refinement as new data becomes available. It starts with an initial training phase, where MD simulations provides sequence scores of ∼60 peptides. This dataset is used to train the ML model, which is used to explore the entire sequence space and predict the fibril-forming probability of new sequences, identifying high-probability candidates. Some of these candidates are then re-evaluated using MD simulations, and the newly obtained data is incorporated into the training set for subsequent iterations. At the end of our last iteration cycle, the 88 peptides account for a training dataset with 22 positive and 50 negative samples–excluding the 16 sequences with MD score of 1.

**Figure 6.**
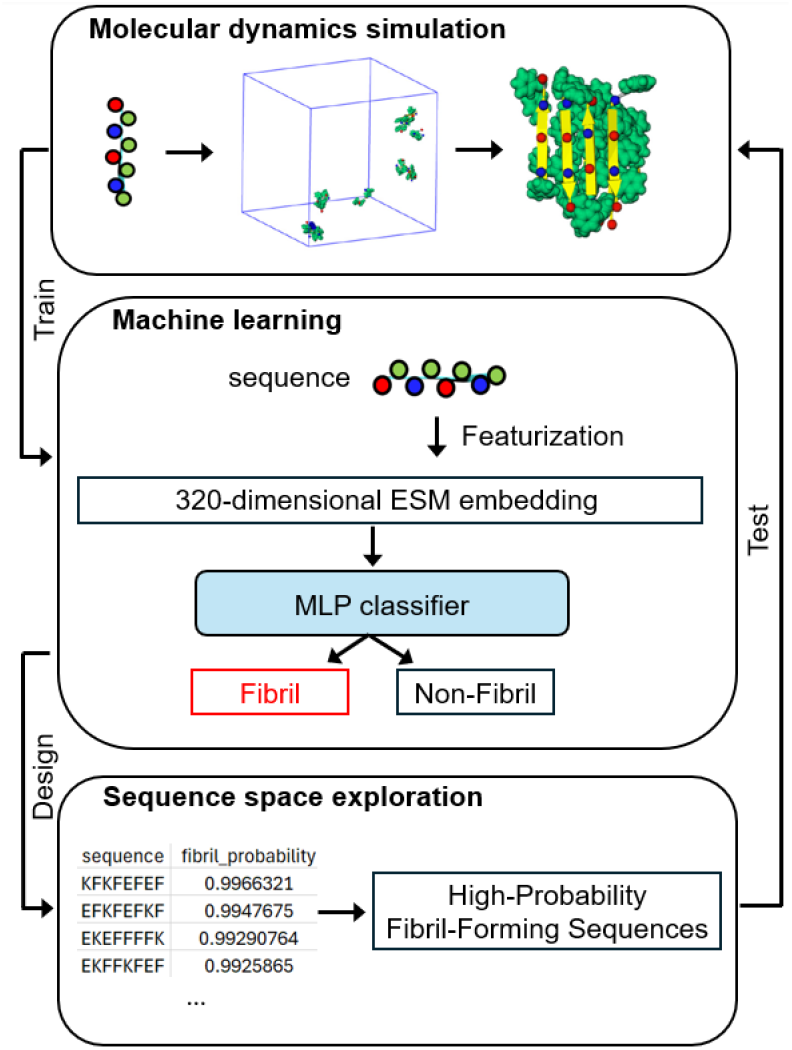
Overview of an active learning framework integrating molecular dynamics (MD) simulations with machine learning (ML) to predict fibril formation in peptides. During the training phase, MD simulations assess the fibril-forming potential of peptide sequences, which are then used to generate labeled data. These sequences are featurized, including a 320-dimensional ESM embedding, and used to train an ML model. The ML classifier (MLP) categorizes each sequence as either “Fibril” or “Non-Fibril.” In the design phase, the trained model is utilized for sequence space exploration to identify high-probability fibril-forming sequences. The newly identified sequences are then validated through MD simulations, creating an iterative loop for continuous model refinement and sequence optimization.

This combination of MD simulations and ML predictions in an active learning loop improves the model in a step-wise manner allowing for an efficient exploration of the sequence space. In particular, by focusing on the most promising fibril-forming candidates allows us to produce a more balanced training set as the overall number of sequences that do not form fibrils formation is much larger. This reduces the computational cost while enhancing the predictive performance and generalizability. Ultimately, the active learning framework enables rapid identification of sequences with high fibril-forming potential, providing a targeted approach to peptide sequence design and evaluation.

## Model Performance

At the end of the last iterative cycle that involved training, prediction, and re-evaluation, the performance of the ML model was computed using 5-fold cross-validation, demonstrating strong performance across several key metrics, as summarized in Table 1. The model achieved an accuracy of 0.944 (± 0.053), indicating consistent and reliable predictions. Precision and recall were both 0.91 (± 0.111), signifying a balanced ability to minimize false positives and capture true fibril-forming sequences. The F1-score was 0.906 (± 0.092), indicating an effective trade-off between precision and recall. The AUC value of 0.939 (± 0.07) further highlights the model’s strong discriminatory power. Overall, these metrics indicate that the model, using only ESM embeddings, has robust predictive capabilities in identifying fibril-forming sequences.

**Table 1:**
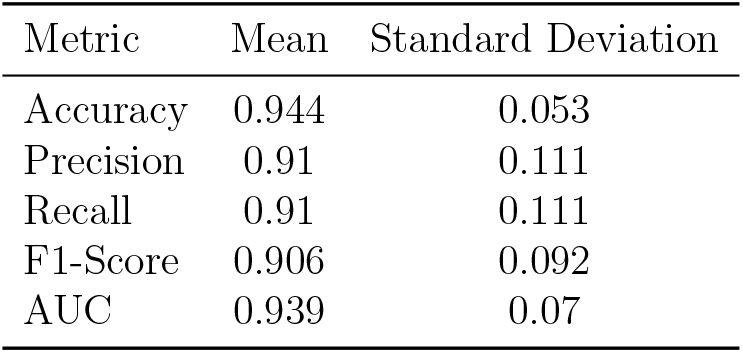
Performance metrics of the ML model evaluated through 5-fold cross-validation.

## ML predictions

Table 2 shows the set of 8 sequences that were simulated to test predictions in the last iterative cycle. In the latter, the model demonstrated a strong capability to predict fibril-forming sequences. Specifically, it successfully identifies four sequences as fibril-forming, all of which had an MD score of 3, indicating high confidence in fibril formation. Additionally, the model accurately classified two sequences as non-fibril-forming, which had an MD score of 0. Interestingly, the sequence FEFEFKFK, highlighted in bold in the table, contains the motif FEFK, which is generally associated with a high propensity for fibril formation. In the training dataset, all sequences containing FEFK formed fibrils. However, our model accurately classified this sequence as non-fibril-forming, consistent with the MD simulation results. This demonstrates the model’s nuanced understanding of fibril formation beyond motif-based assumptions.

**Table 2:**
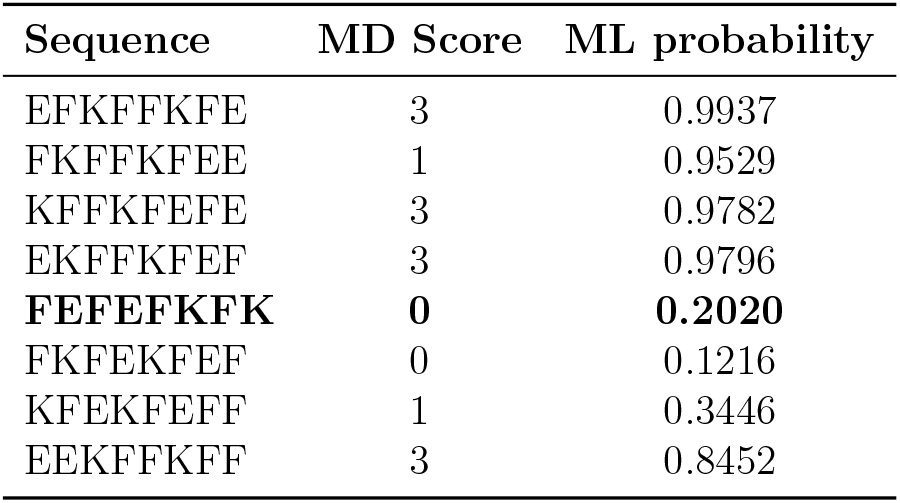
Comparison between MD simulation scores and ML predicted probabilities for fibril formation of peptide sequences. The “MD Score” indicates the number of MD simulations where fibril formation was observed (range 0–4), and “ML Probability” is the probability of fibril formation predicted by the machine learning model.

Fig. 7 shows the probability of forming fibrils for sequences that were not simulated but exhibit the same polar-nonpolar sequence patterns as in Fig. 4. This violin plot confirms that sequences with pattern `FF__FF` (orange) and `F__FFF` (red) are not likely to form fibrils. Conversely, almost all sequences where non-polar and polar residues alternate (`F_F_F_F_` and `_F_F_F_F` in black and green, respectively) tend to form fibrils. Notice that in Fig. 4, the reduced set of simulated sequence with pattern `FFF_F` (brown) showed a high probability to form fibrils as opposed to its reversed pattern `F_FFF` (yellow) that inhibited fibril formation. Fig. 4 shows that when all sequences of the reversed `F_FFF` pattern are considered, the probability of forming fibrils covers a broad spectrum ranging from zero to one. This highlights the danger of drawing conclusions based on results from a small set of simulations/experiments. The grouping of all nonpolar residues next to each other, i.e., FFFF (blue), also accounts for a broad probability spectrum with many sequences predicted to form fibrils.

**Figure 7.**
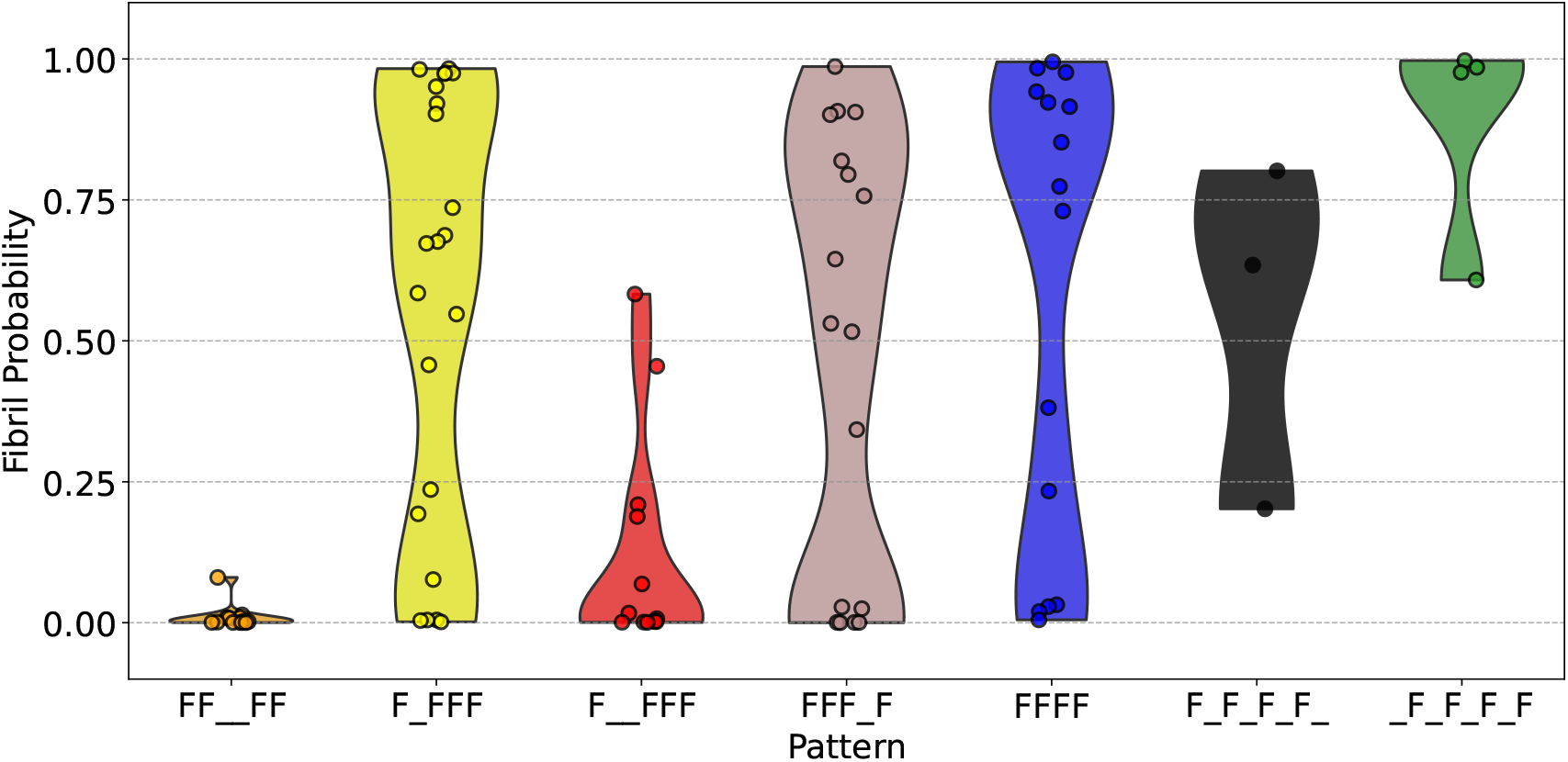
Violin plot showing the distribution of fibril-forming probabilities for sequences with known patterns.

Fig. 8 shows the probability of forming fibrils for all polar-nonpolar patterns excluding the ones shown in Fig. 7. Only sequences in 5 patterns form fibrils with a probability higher than 0.5. Two of these patterns, i.e., `F_FF_F` (red) and `FF_FF` (purple), account for peptides with only two non-polar residues facing the same direction when adopting an extended conformation. The other three patterns exhibit three non-polar residues alternating with a charged amino acid (blue, green, and orange). All other patterns do not show a significant probability to form fibrils.

**Figure 8.**
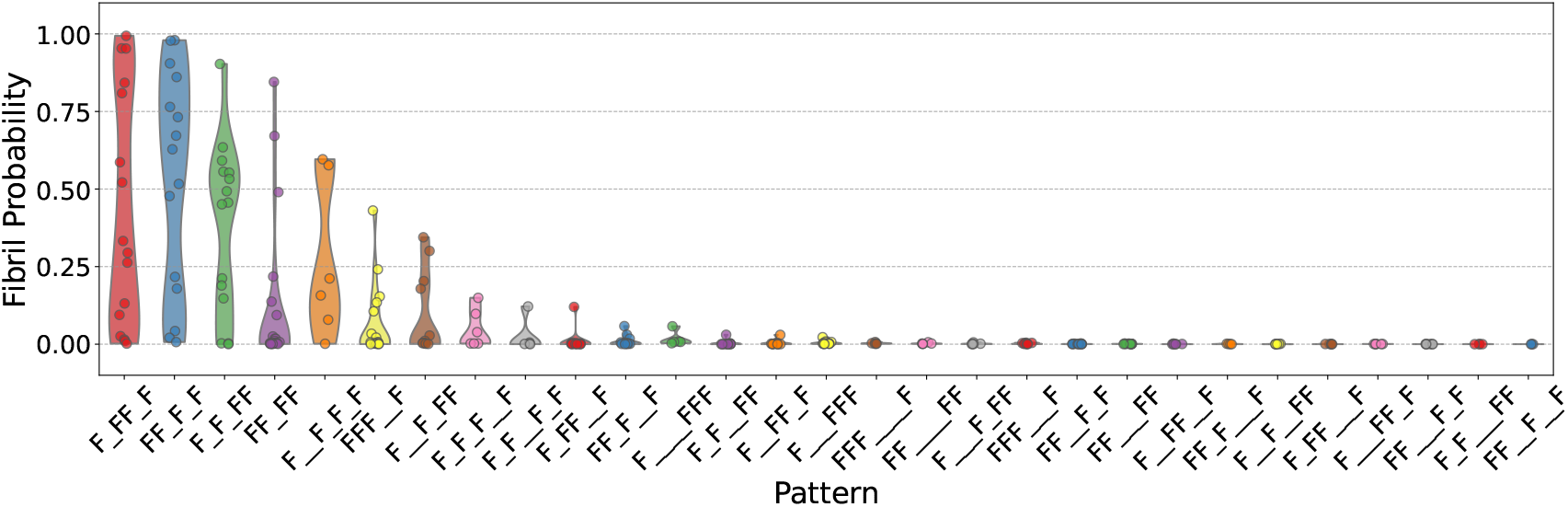
Violin plot showing the distribution of fibril-forming probabilities for sequences with the other patterns.

To summarize the results of the ML model, we show a graph with the set of ten (out of a total of 35) patterns that exhibit at least one sequence that forms fibrils in Fig. 9. In the latter, `F_F_F_F_` and `_F_F_F_F` are grouped within the same `F_F_F_F` pattern. The 10 patterns are displayed (panel b) from left to right in terms of the maximum number of non-polar residues that face the same direction when peptides are folded in an *β*-strand. For clarity, patterns are connected by a line if they can be transformed into each other by swapping any non-polar residue *i* with one of its neighbors *i* + 1 or *i* − 1. This corresponds to performing mutations that preserve the number of non-polar residues in the sequence. Accordingly, counting the number of intermediate patterns needed to connect two sequences provides a measure of their proximity. For example, transforming `F_F_F_F` into `FFFF` requires swapping phenylalanine at least four times with a polar residue. In this graph, symmetry is highlighted by patterns that have many sequences that form fibrils (blue circles in the figure). They are either palindromes (`F_F_F_F, FFFF`, and `F_FF_F`) or their reversed sequence also forms fibrils (`F_FFF` with `FFF_F`, and `F_F_FF` with `FF_F_F`). These symmetric sequences are interconnected to each other accounting for a small and tight graph of fibril forming patterns.

**Figure 9.**
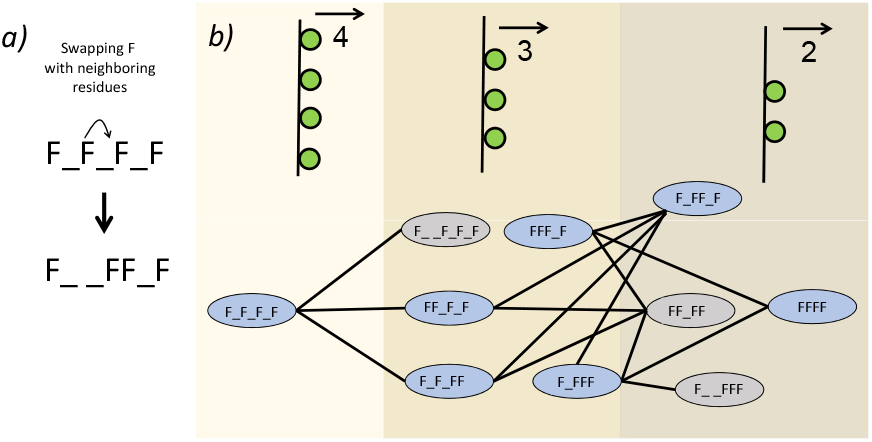
Graph of the 10 fibril-forming patterns identified by the ML model. (a) Two non-polar patterns are connected by a line in the graph if they can be transformed into each other by swapping one of their phenylalanines with a neighboring residue. (b) Non-polar patterns with 4, 3, and 2 non-polar residues facing the same direction when folded into a *β*-strand are displayed from left to right. Patterns with many fibril-forming sequences are colored in blue.

## Discussion and Conclusion

In this study, we employed an active-learning framework to investigate how specific peptide sequence properties govern fibril-forming propensities. By integrating unbiased molecular dynamics (MD) simulations with machine learning (ML), we harnessed the high-resolution insight of atomistic simulations in concert with the efficiency and predictive capabilities of data-driven artificial intelligence models. This synergy enabled us to accurately predict fibril formation across a wide array of peptide sequences, demonstrating the efficacy of combining physics-based simulations with ML classification methods.

Our MD simulation results demonstrate that even subtle variations in residue patterning, and charge/hydrophobic distribution can strongly affect the nucleation rate of amyloid-like fibrils. In particular, sequence-specific interactions drive the formation of cross-*β* architectures, underscoring the complexities of peptide self-assembly. These insights deepen our understanding of how discrete sequence changes modulate protein misfolding and aggregation, offering valuable design principles for peptide-based materials. We further show that short, high-concentration simulations in small boxes can effectively serve as a proxy for longer, large-box simulations during data curation. Overall, this work offers a systematic strategy for both rapid assessment and rational design of amyloid-forming peptides. By unifying MD simulations and ML approaches within an active-learning loop, we present a versatile framework that may expedite the discovery of fibril-prone sequences and guide the engineering of functional amyloid-like systems. We anticipate that these findings will be of broad interest for applications in biomaterials and therapeutics, where precise control over peptide aggregation is essential.

## Methodology

### Molecular Dynamics Simulations

All-atom molecular dynamics simulations were performed using the GROMACS 2020 package^50^ with the Amber99sb-ILDN force field ^51^for proteins and the TIP3P water model^52,53^ for solvation. Systems were set up in cubic boxes with periodic boundary conditions and simulated in the isothermal–isobaric (NPT) ensemble. Pressure was isotropically maintained at 1 bar using the Parrinello–Rahman barostat^54^ (*τ*_*p*_ = 2.0 ps), while temperature was controlled at 350 K by coupling the solute and solvent separately to a velocity-rescaling thermostat^55^ (*τ*_*t*_ = 0.1 ps). The equations of motion were integrated using the leapfrog algorithm with a 2 fs time step. Short-range van der Waals and electrostatic interactions were truncated at 1.0 nm, and long-range electrostatic interactions were computed via the smooth Particle Mesh Ewald (PME) method.^56,57^ Prior to production runs, each system underwent sequential equilibration in both the NVT and NPT ensembles for a total of 100 ps. Simulation details specific are described in the Results sections.

### Analysis of MD Trajectories

The trajectories generated in our MD simulations were analyzed using a combination of built-in GROMACS tools and custom in-house scripts. Key metrics were extracted to characterize peptide aggregation, including the minimum distances between backbone atoms of different peptides and the secondary structure content determined by the DSSP algorithm.^58^ In addition, GROMACS was used to identify all peptide pairs that came within 0.5 nm of each other; this list served as input for our custom clustering code, ^12^ which calculates the size of the largest aggregate (or cluster) formed by peptides in contact. The number of peptides in the largest cluster was used as a measure of nucleus formation. Furthermore, hydrogen bonding was monitored by evaluating the geometric criteria for hydrogen bonds: a donor-acceptor distance of ≤0.35 nm and an H–D–A angle of ≤ 30°. These analyses enabled us to quantify both the extent of secondary structure formation and the evolution of aggregate size.

### Performance of the ML model

To evaluate the performance of the final model after several iterations of active learning, we employed 5-fold cross-validation during the final iteration. This approach involves splitting the dataset into five subsets, using each subset once as a validation set while training the model on the remaining subsets. This method ensures robust model performance and reduces the risk of overfitting to any particular subset of the data. The model’s performance was evaluated using several key metrics:

#### Accuracy

This metric represents the proportion of total predictions (both fibril and non-fibril) that the model classified correctly. It serves as a general indicator of model performance but may be less informative when dealing with imbalanced datasets. The formula for accuracy is given by:

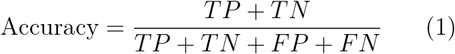

Where:

- TP: Correctly predicted fibrils.
- TN: Correctly predicted non-fibrils.
- FP: Incorrectly predicted fibrils.
- FN: Incorrectly predicted non-fibrils.

#### Precision

Precision, also known as the positive predictive value, is the proportion of true positive predictions (fibril) out of all positive predictions made by the model. It is given by the formula:

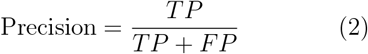

High precision indicates that when the model predicts a sequence to form fibrils, it is often correct.

#### Recall

Also known as sensitivity or true positive rate, recall measures the proportion of actual fibril-forming sequences that are correctly identified by the model. High recall ensures that the model is effective at finding most of the fibril-forming sequences, which is crucial in screening applications where false negatives are costly. Mathematically, recall is given by:

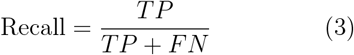

#### F1-Score

The F1-score is the harmonic mean of precision and recall. It provides a balance between these two metrics, especially useful when we need to find an equilibrium between identifying fibril-forming sequences while minimizing false positives. It is expressed as:

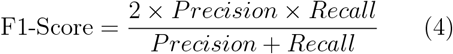

### Area Under the Receiver Operating Characteristic Curve (AUC)

The AUC measures the model’s ability to distinguish between fibril and non-fibril sequences across all classification thresholds. An AUC close to 1 indicates strong discriminatory power, meaning the model is effective at distinguishing positive from negative cases.

## Data and Code Availability

All processed datasets and the active learning workflow are available on GitHub at Fibril Prediction

## Acknowledgements

Z.S. acknowledges support from the Vanderbilt Data Science Postdoctoral Fellowship. X.K. gratefully acknowledges the research funding and support provided by the Vanderbilt Data Science Institute. C.D. and S.J. were supported by the National Institute of General Medical Health under grant no. 1R15GM148982-Computational resources were provided by the Academic and Research Computing System (ARCS). J.W. acknowledges the supports from the National Science Research and Innovation fund of Thailand (NSRF) via the Program Management Unit for Human Resources and Institutional Development Research and Innovation (PMUB) [grant No. B42G670041], the National Research Council of Thailand (NRCT) through the Research Grants for Talented MidCareer Researchers (grant no. N41A640080), and the Kasetsart University Research and Development Institute (KURDI) via the Fundamental Fund (grant no. FF(KU)33.67).

